# Enhancing coevolution-based contact prediction by imposing structural self-consistency of the contacts

**DOI:** 10.1101/279190

**Authors:** Maher M. Kassem, Lars B. Christoffersen, Andrea Cavalli, Kresten Lindorff-Larsen

## Abstract

Based on the development of new algorithms and growth of sequence databases, it has recently become possible to build robust and informative higher-order statistical sequence models based on large sets of aligned protein sequences. By disentangling direct and indirect effects, such models have proven useful to assess phenotypic landscapes, determine protein-protein interaction sites, and in *de novo* structure prediction. In the context of structure prediction, the sequence models are used to find pairs of residues that co-vary during evolution, and hence are likely to be in spatial proximity in the functional native protein. The accuracy of these algorithms, however, drop dramatically when the number of sequences in the alignment is small, and thus the highest ranking pairs may include a substantial number of false positive predictions. We have developed a method that we termed CE-YAPP (CoEvolution-YAPP), that is based on YAPP (Yet Another Peak Processor), which has been shown to solve a similar problem in NMR spectroscopy. By simultaneously performing structure prediction and contact assignment, CE-YAPP uses structural self-consistency as a filter to remove false positive contacts. At the same time CE-YAPP solves another problem, namely how many contacts to choose from the ordered list of covarying amino acid pairs. Our results show that CE-YAPP consistently and substantially improves contact prediction from multiple sequence alignments, in particular for proteins that are difficult targets. We further show that CE-YAPP can be integrated with many different contact prediction methods, and thus will benefit also from improvements in algorithms for sequence analyses. Finally, we show that the structures determined from CE-YAPP are also in better agreement with those determined using traditional methods in structural biology.

**Author summary:** Homologous proteins generally have similar functions and three-dimensional structures. This in turn means that it is possible to extract structural information from a detailed analysis of a multiple sequence alignment of a protein sequence. In particular, it has been shown that global statistical analyses of such sequence alignments allows one to find pairs of residues that have covaried during evolution, and that such pairs are likely to be in close contact in the folded protein structure. Although these insights have led to important developments in our ability to predict protein structures, these methods generally result in many false positive contacts predicted when the number of homologous sequences is not large. To deal with this issue, we have developed CE-YAPP, a method that can take a noisy set of predicted contacts as input and robustly detect many incorrectly predicted contacts within these. More specifically, our method performs simultaneous structure prediction and contact assignment so as to use structural self-consistency as a filter for erroneous predictions. In this way, CE-YAPP improves contact and structure predictions, and thus advances our ability to extract structural information from analyses of the evolutionary record of a protein.

## Introduction

A large and recent increase in known protein sequences has sparked an interest in using the multiple sequence alignments (MSAs) of protein families to predict native contacts in globular proteins [1], membrane proteins [2,3], as well as predicting contacts in protein-protein interfaces [4,5] Conceptually speaking, homologous proteins from diverse organisms are likely to have similar 3D structures due to the conservation of function [6]. As a result, the sequence space explored across a protein family is highly constrained. Of special interest, are pairwise coevolving amino acid positions in the MSA. These coevolution patterns have been shown to correlate strongly with spatial proximity in the native 3D structure [7].

Methods initially used to quantify the degree of pairwise positional coevolution were based on local statistical models (e.g. mutual information) that do not disentangle transitive effects often seen in proteins. Current state-of-the-art methods rely on global statistical models (e.g. maximum entropy), well-known from statistical physics, to help disentangle transitive effects and, thereby, provide more robust and precise contact predictions. The maximum entropy principle is increasingly used in computational biology because of its ability to produce accurate global models given observed data (e.g. an MSA) with minimal risk of overfitting [8]. The apparently first to use the maximum entropy principle in the coevolution analysis of protein sequences was Lapedes et al. [9]. Similar but more recent methods are the mean field Direct Coupling Analysis approach [1, 4, 10] followed e.g. by pseudo-likelihood maximization [11,12], sparse inverse covariance estimation approach [13], and various machine learning methods [14–16]. Many of the methods have recently been reviewed extensively [17].

An obvious and popular use of predicted contacts is to implement them as distance restraints in protein folding simulations. The restraints can dramatically reduce the conformational search space, thereby enabling structure calculations of even large (> 250 amino acid residue) proteins [1]. One major challenge is, however, that the number of effective sequences (definition in Methods) needs to be sufficiently large (>∽ 5 times the number of amino acids [18]) to ensure that enough contacts can be predicted accurately. Protein families with a sufficiently large number of sequences are, however, also more likely to have at least one experimentally solved structure, which, makes template-based modeling a more viable option [18]. A key challenge is, therefore, to decrease the required number of effective sequences to a level that enables the precise contact predictions of more protein families without experimental structures. Recently, the number of protein families, with a sufficient number of effective sequences and without homologous structures, was estimated to be ∽400 [18]. To increase this number and thereby decrease the required number of effective sequences, developers attempt to improve (or combine [19]) the statistical models. While there might be room for improvement, it is possible that these methods will not reach the level of precision needed to consistently compute accurate protein structures without a significant number of homologous sequences. There are, however, examples of experimentally difficult protein targets without solved structures that had enough sequences to solve the structures using coevolution [2,3,20,22] –.

There are two initial obstacles to overcome when using predicted contacts in structure prediction. First, one needs to decide how many contacts to include. The methods described above simply rank contacts by decreasing strength of the coevolutionary signal, but does not directly provide a natural cutoff for how strong the signal should be in order to consider a pair of residues likely to be in contact. Secondly, even with many sequences and conservative choices for how many contacts to use, one generally ends up with a number of false positive (FP) predictions, i.e. pairs of residues that show some level of coevolution, but are not in close proximity in the three-dimensional structure. In practical applications, these two problems are tightly related: One would like to include as many contacts as possible to restrain the three dimensional structure, but at the same time risk including many FPs. For example, one would on average expect ∽5 of the top 20 (i.e. 25%) coevolving pairs of residues to be FPs for a 100-residue long protein with an MSA with 500 sequences, increasing to ∽ 20 of the top 50 (40%) coevolving pairs to be FPs [18].

To circumvent the problem of noisy predictions, MacCallum and co-workers have suggested an elegant approach termed MELD [23]. The basic idea in MELD is to explicitly take into account that a fraction of the predicted contacts are wrong, and hence should not be included. In practice this is done by iteratively dividing contacts into either an ‘active’ or ‘ignored’ set, with those contacts that agree the worst with the current structural model partitioned into the ignored fraction. Thus, using the example from above, if we know that ∽ 20 of the top 50 contacts are FPs, but not which of them, we only consider as active those 30 contacts that agree best with the structure. In this way structural self-consistency is used to guide contact assignment and structure prediction at the same time. One key limitation of this approach is that it is not always clear how large a fraction of the contacts can be ignored. Even if we know the fraction of FPs that will be present on average, it is difficult to predict this number specifically for a given protein. A different approach is to include experimental data, such as from NMR, and use self-consistency as a filter to remove false positives [24].

Here, we describe a method, called CE-YAPP, which we have developed to simultaneously determine the number of long range predicted co-evolution contacts (PCCs) and to partition these contacts into true positives (TPs) and FPs (Fig. 1). The method builds upon the automated nuclear magnetic resonance (NMR) NOESY structure determination method called YAPP (Yet Another Peak Processor) [25].

**Fig 1.**
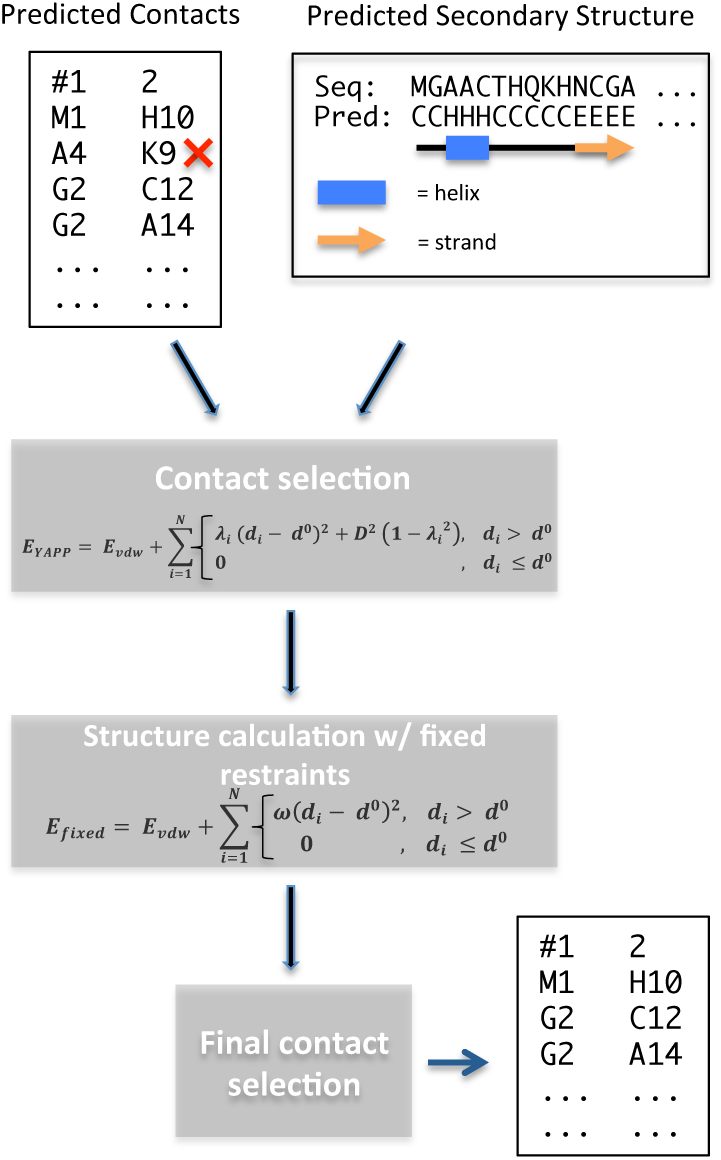
Workflow diagram of the CE-YAPP method. Predicted coevolution contacts and predicted secondary structure are used in combination to filter out false positive contacts. The red ‘x’ represents a false positive contact.

YAPP automatically assigns NOESY peaks to infer distance restraints that are subsequently used in a structure calculation. In NMR, these restraints are often the only source of information used to determine protein structures. In contrast, we designed CE-YAPP to use long-range PCCs as distance restraints and combine them with local structural information in the form of secondary structure predictions. Both YAPP and CE-YAPP share a unique protocol in which distance restraints that are in systematic violation of the protein geometry during structure calculations are automatically detected and turned off. The false-positive-detection is carried out by sampling, for each individual distance restraint, a parameter that allows us to turn off this contact with some energetic cost. These contacts/restraints are then identified as likely FPs and are discarded from the initial list of predicted contacts, thus, enhancing the contact precision by keeping the TPs.

We tested CE-YAPP using a recently-described data set, called Noumenon, which consists of 150 MSAs and their associated crystal structures [26]. This data set has been curated to remove the selection bias seen when using protein families that have at least one experimentally solved structure. Our results show that CE-YAPP provides an effective solution to the problem of both finding a useful number of contacts and filtering FPs in a noisy prediction. In particular, we find that CE-YAPP increases prediction accuracy also when there are fewer number of sequences available. We also show that CE-YAPP can be combined with different methods for contact prediction, suggesting that the approach can be used generally to improve predictions even as methods for contact prediction continue to improve.

## Results and Discussion

### A framework for detecting structural self-consistency

The main goal of this work was to develop a method that enhances the precision of PCCs. We developed CE-YAPP which achieves this goal by taking an automatically chosen set of long-range (sequence-wise) PCCs and identifying the FP contacts within these. CE-YAPP performs simulations that incorporate predicted secondary structure and makes geometrical considerations of each PCC to remove those that are systematically inconsistent with the geometry (Fig. 1). More specifically, CE-YAPP begins by building an extended protein structure with fixed ideal secondary structure segments (ideal α-helix or β-strand), based on the predicted secondary structure. The segments are structurally defined using ideal ϕ and ψ dihedral angles for the residues predicted to be α-helical or β-stranded. Subsequently, CE-YAPP performs rapid simulations, using the chosen subset of PCCs as distance restraints and allows only changes to the dihedral angles that are not fixed. A computationally-efficient energy function, that includes a van der Waals term and a restraints term, controls the structure calculation while automatically identifying systematically violated distance restraints, by sampling the weights, *>* _*i*_ (see Methods).

A general issue when using PCCs for structure calculation, is the need to decide the number of PCCs to use. The issue becomes especially problematic when there are only a few effective sequences (e.g. *<* 5 sequences per *N*_*AA*_; the number of amino acids) available, due to the higher risk of observing FP contacts [18]. In CE-YAPP we solve this issue by including a relatively large number of contacts, but then effectively filter away the FPs through requiring structural self-consistency. Specifically, we include 1.2 *N*_*AA*_ contacts between pairs of residues that are not both within a single predicted secondary structure segment (i.e. are long-range). The algorithm is robust to the exact choice of the number of contacts included (see Supporting Information for additional details).

To illustrate the idea and performance of CE-YAPP, we show the results for the 95 amino acid residues long *E. coli* ribosome hibernation promoting factor (PDB ID: 2RQL), using ∽ 600 effectives sequences for the contact prediction (Fig. 2). In this specific case, the number of input contacts was 114 (*N*_*input*_ = *N*_*AA*_ 1.2 = 114). Thus, we first sort contacts by the strength of the evolutionary couplings and find the top 114 contacts that are not within a single predicted secondary structure element. Comparison with the known NMR structure reveals that 82 of these are TPs corresponding to a precision of 72 %. To increase the precision, CE-YAPP repeats the simulation protocol (Fig. 1) 64 times and discards contacts that are turned off in more than 70 % of the runs (Fig. 2A). In doing so, CE-YAPP retains 102 of the 114 contacts (CE-YAPP contacts) reducing the number of FP contacts from 32 to 21, thereby, increasing the precision from 72 % to 79 %. These results can be visualised in the context of the experimental contact map (Fig. 2A) which shows how most of the contacts excluded by CE-YAPP correspond to FPs, demonstrating the power of the approach in identifying a self-consistent set of contacts. The map also reveals several apparently FP contacts that are not removed by CE-YAPP. It is clear, however, that many of these are close (in sequence) to true contacts, and many of them are just outside the distance range that we use to define contacts. Thus, we conclude from this example (i) that CE-YAPP has the potential to identify a number of self-consistent contacts from a list of noisy contacts, (ii) that the algorithm can remove many FPs with only minimal loss of TPs and (iii) it appears that at least some of the FPs that are not removed by CE-YAPP are only ‘borderline errors‘.

**Fig 2.**
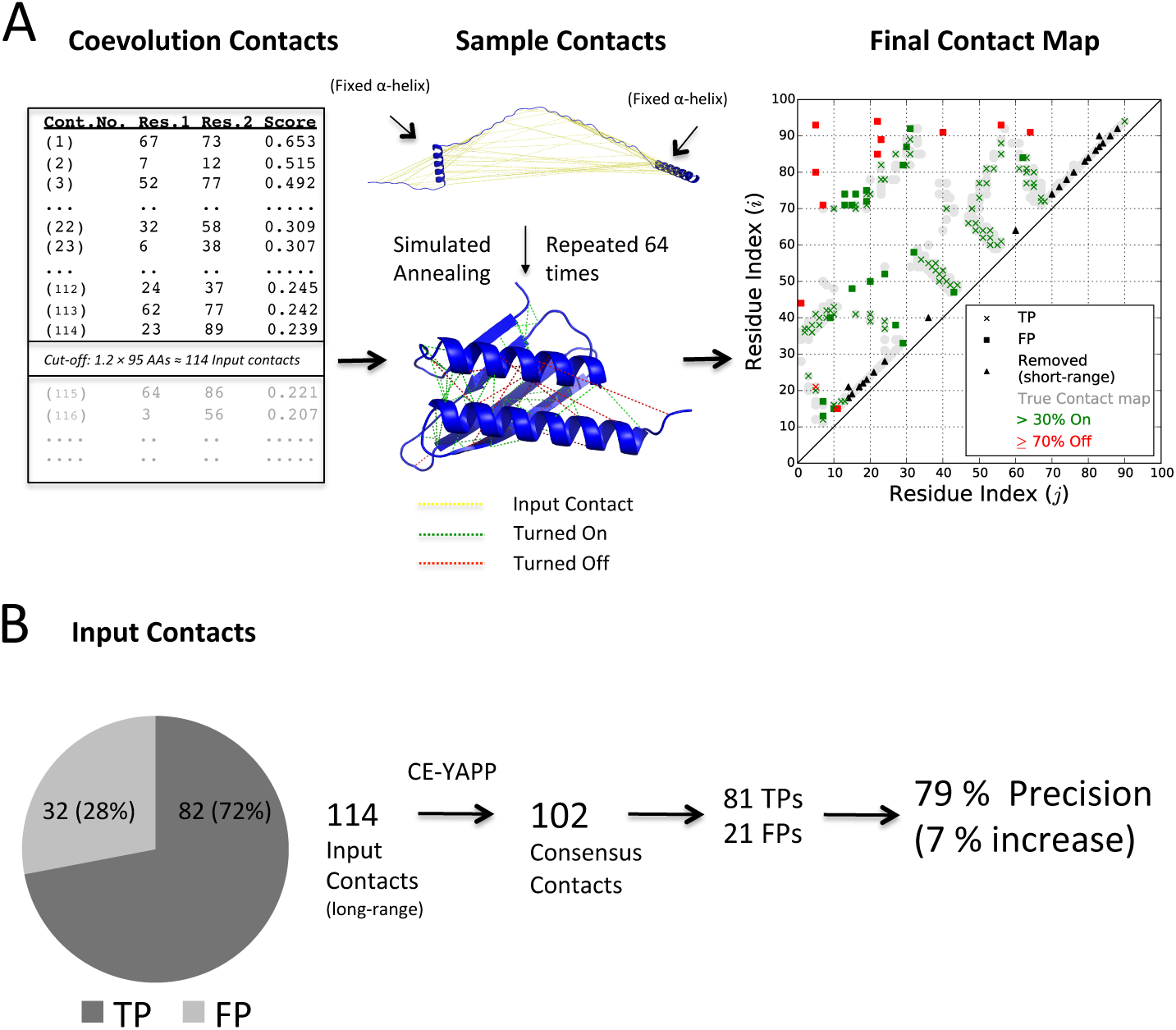
CE-YAPP Protocol and results for the ribosome hibernation promoting factor HPF. (A) CE-YAPP uses as input 114 coevolution based long-range contacts predicted using Gremlin [11]. These contacts are then used as input to the protocol depicted in Fig. 1, and repeated 64 times producing 64 similar contact lists. The final list of predicted consensus contacts are those that are turned on in more than 30 % of the simulations. (B) The precision of the consensus contacts produced by CE-YAPP is compared to the precision of the input set of contacts.

### Benchmarking CE-YAPP

Encouraged by these initial observations, we continued to benchmark the performance of CE-YAPP using several indicators such as precision, recall, and number of contacts. In these analyses we used the recently described Noumenon data set [26], which contains 150 proteins with known structures and a representative distribution of sequence depths (i.e. effective sequences); in practice we performed our analysis of 147 of these proteins (see Methods). The results, summarised in Fig. 3, demonstrate that CE-YAPP consistently improve the accuracy of contact predictions. The proteins have been sorted according to the depths of their MSAs, quantified as the number of effective sequences divided by the number of amino acids (*N*_*Eff*_ */N*_*AA*_; Fig. 3A). The number of input contacts (fixed at 1.2 *N*_*AA*_) and the number of contacts after running CE-YAPP are shown in Fig. 3B. As expected, when there are many effective sequences,CE-YAPP discards only few contacts whereas many are filtered away when there is only little information in the MSA. This behaviour can be rationalised given that coevolution-based contact predictors (e.g. Gremlin [11]) generally produce contacts with lower precision when *N*_*Eff*_ */N*_*AA*_ is low, prompting CE-YAPP to discard more contacts. We also calculated the recall, i.e. the fraction of TPs in the contact list that are retained after CE-YAPP filtering (Fig. 3C). At high *N*_*Eff*_ */N*_*AA*_ (*>* 5), the recall is close to one, meaning that CE-YAPP rarely discards TP contacts in this region. In the intermediate region (1 *< N*_*Eff*_ */N*_*AA*_ < 5) the recall is ∽0.8 – 0.9.

**Fig 3.**
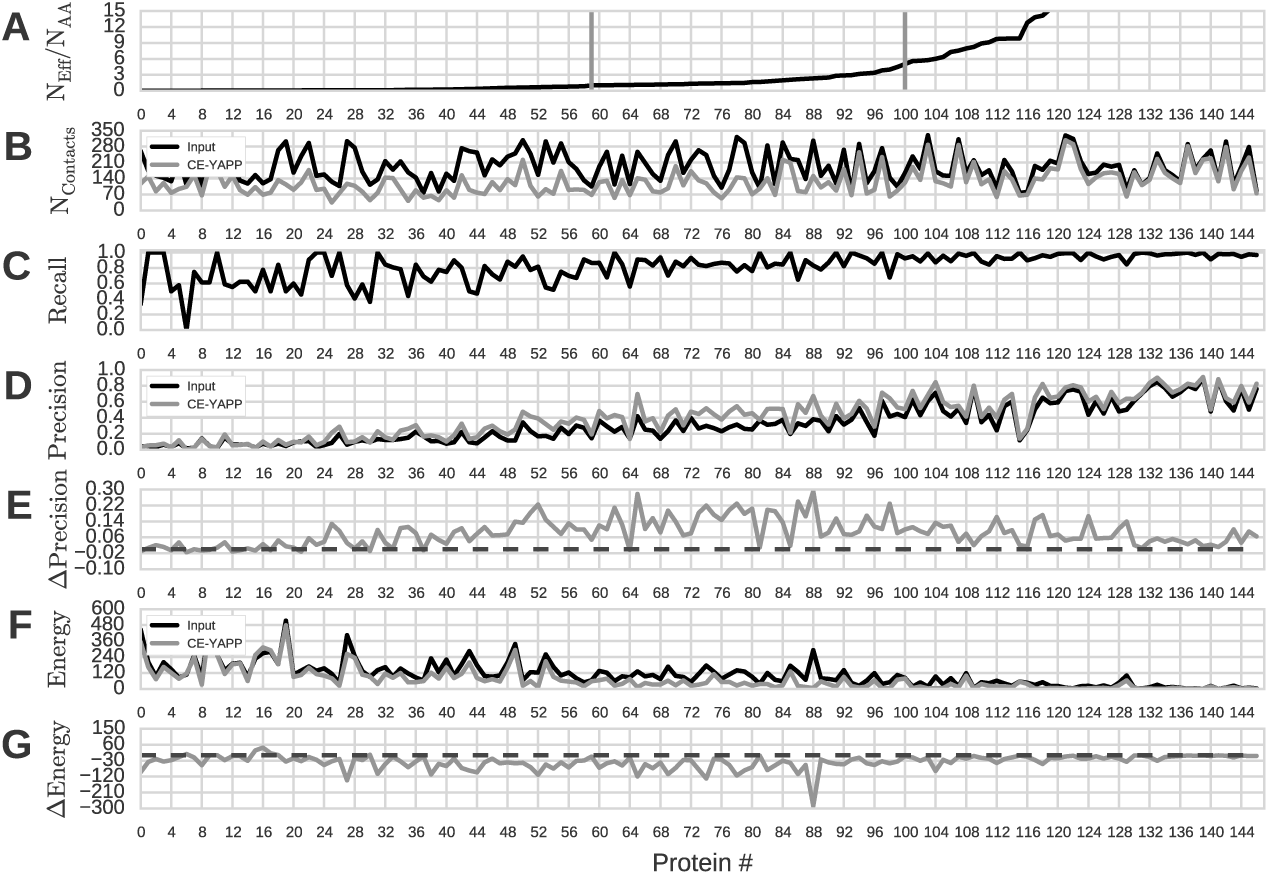
CE-YAPP performance on the Noumenon dataset. A: The number of effective sequences divided by the number of amino acids, *N*_*Eff*_ */N*_*AA*_, is plotted for each protein and sorted from low to high. The data in the remaining panels are sorted accordingly. The grey vertical bars represent the proteins with *N*_*Eff*_ */N*_*AA*_ closest to 1 and 5, respectively. B: Number of contacts. C: Recall (*TP/*(*TP* + *FN*)) of the CE-YAPP contacts. D: Precision (*TP/*(*TP* + *FP*)). E: Precision of CE-YAPP contacts minus precision of the input contacts (Δ. *Precision*). The black dashed lines in panels E and F denote zero. F: Restraint violation energy (Eq. 4) G: Drop in restraint violation after CE-YAPP (Δ. *Energy*).

These results are encouraging as they suggest that CE-YAPP, even with only modest amounts of sequences, can find a consistent set of contacts that contain most of the TPs in the input set. An equally if not more important measure of performance is precision, which quantifies the fraction of contacts that are TPs. Comparison of the precision in the input contacts and the output from CE-YAPP shows a consistent improvement in precision, i.e. that CE-YAPP is able to filter away FP contacts. Again, as expected, precision is greatest at high values of *N*_*Eff*_ */N*_*AA*_ and drops as the information content in the MSA decreases (Fig. 3D). It is clear, however, that there is a general increase in precision after CE-YAPP filtering (Fig. 3E, which shows the increase in precision after CE-YAPP). This improvement is especially pronounced with values of *N*_*Eff*_ */N*_*AA*_ in the range 1–5 — a region that generally includes protein families that contact-based structure prediction find to be too difficult [18]. The average improvement in precision is 0.07 for *N*_*Eff*_ */N*_*AA*_ *>* 5 and 0.14 for 1 *< N*_*Eff*_ */N*_*AA*_ < 5.

As discussed above in the example with the ribosome hibernation promoting factor HPF (Fig. 2) we observed that the FPs, that CE-YAPP did not remove, appeared to be close to real contacts. Because precision does not quantify the severity by which FPs violate the TP definition, we also calculated the weighted mean squared distance violations (’energy‘; Eq. 4) of the contacts with respect to the PDB structures (Fig. 3F), and the change of these violations after CE-YAPP (Fig. 3G). Similar to our observations using precision, we find that CE-YAPP improves contact prediction also when judged by restraint violations, and that the improvement is large also in the region with intermediate values of *N*_*Eff*_ */N*_*AA*_ (Fig. 3G).

As expected, we note that when there are many sequences (*N*_*Eff*_ */N*_*AA*_ *>* 5), the energy of the input contacts is significantly lower than when there are an intermediate or low number of sequences (Fig. 3F). Interestingly, this can be the case even when the apparent precision is low. Examples of this behaviour is observed for protein number 112 and 115, where the precision of the input contacts is low (∽20%) but with energies close to zero. This suggests that the predicted contacts are close to the boundary between TPs and FPs, albeit more often on the ‘FP side‘, highlighting an issue regarding precision as a performance measure.

### Improved structural accuracy

Together, the results described above demonstrate how CE-YAPP can be used to find a self-consistent set of contacts, and how this algorithm is able to increase precision in contact prediction. One application of contact prediction is in protein structure prediction, where contact-assisted protein folding has enabled new progress in our ability to predict protein structure from amino acid sequence(s). Thus, we set out to examine whether the improved contact prediction also translates into improved quality of three dimensional structures. In these calculations we continued to work with the long-range contacts that are the focus on CE-YAPP, but in contrast to the work described above we decided to use the actual backbone dihedral angles in the secondary structures of the experimentally-derived structures. We thus determined the *ϕ* and dihedral angles of ordered secondary structure regions from the respective PDB structures and fixed these dihedral angles to those values. This ensures that the secondary structure of our structure calculations matches exactly the secondary structure of the experimental structures such that we can pin down the effect of the contacts on the tertiary structure. For the same reasons, we refrained from using a complex force field to give a better picture of the contribution of the contacts to the structures, and thus used only a restraint potential and a van der Waals excluded volume term. As a control for the maximum performance possible, we also performed calculations using only the TP contacts from within the top-ranking contacts.

We performed 16 structure calculations for each protein and for each of the three contact sets (only TPs, before and after CE-YAPP filtering), and report the average across those repetitions, again sorting the proteins according to *N*_*Eff*_ */N*_*AA*_ (Fig. 4). As a measure of structural quality, we chose the global distance test (GDT) with a single cutofcves indicating good agreement between calculated and experimental structures (Fig. 4B). Not surprisingly, we observe that the true contact sets generally outperforms both the input contacts and the CE-YAPP contacts, and with average GDT(5) values of 0.22, 0.60 and 0.73 in the three ranges of *N*_*Eff*_ */N*_*AA*_(*<* 1, 1–5 and *>* 5, respectively). The high values obtained when *N*_*Eff*_ */N*_*AA*_ *>* 1 suggests that there are sufficiently many real contacts among the top 1.2 *N*_*AA*_contacts to determine a reasonably accurate structure of the proteins.

**Fig 4.**
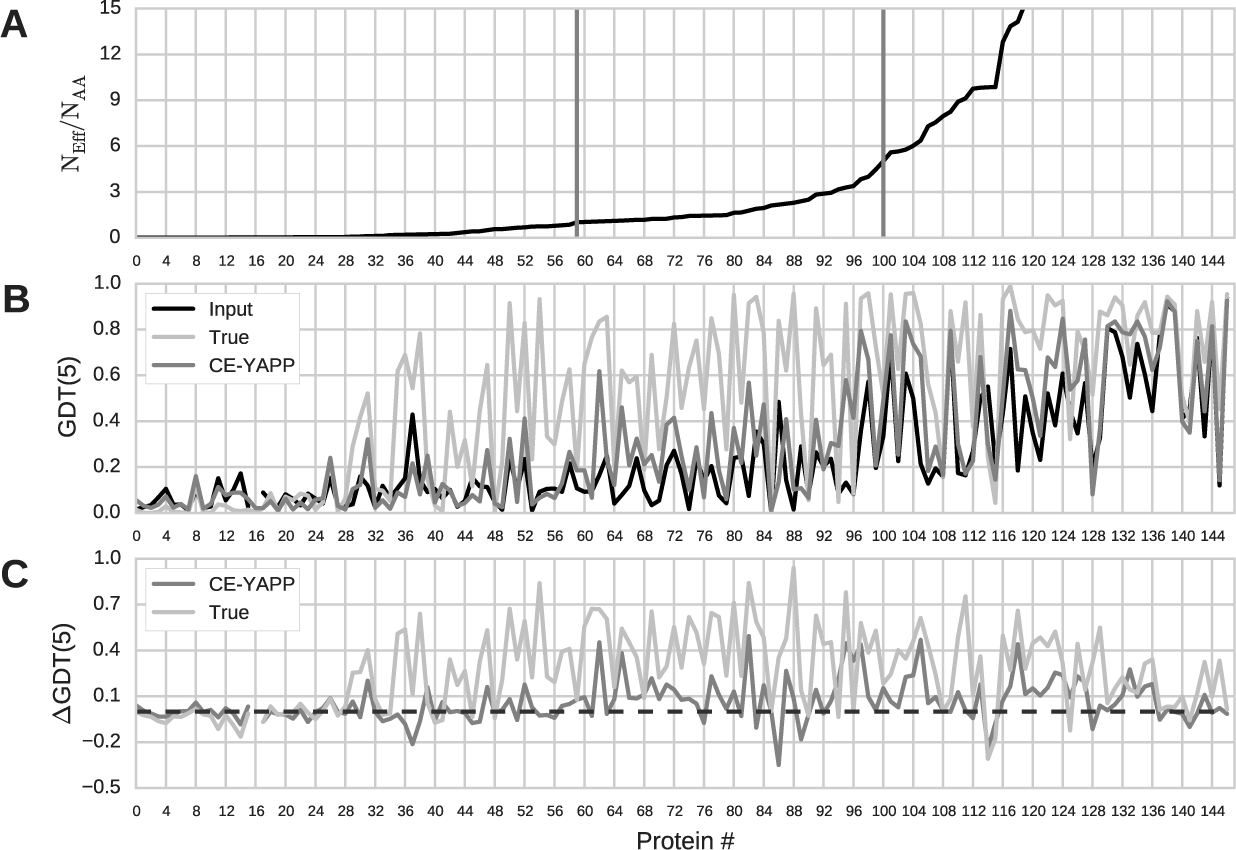
Structural Performance on the Noumenon dataset. Panel 1: The number of effective sequences divided by the number of amino acids, *N*_*Eff*_ */N*_*AA*_, is plotted for each protein and sorted from low to high. The data in the remaining panels are sorted accordingly. The grey vertical bars represent the proteins with *N*_*Eff*_ */N*_*AA*_ closest to 1 and 5, respectively. Panel 2: GDT with a single cutoff of 5 ^A°^. Panel 3: Difference in GDT(5) (Δ*GDT* (5)). The black dashed line denotes zero.

In the same three ranges of *N*_*Eff*_ */N*_*AA*_ (*<* 1, 1–5 and *>* 5) the average values of GDT(5) are 0.09, 0.17 and 0.48 when using contacts before CE-YAPP filtering and 0.09,and 0.57 after filtering. From these results we make two observations. First, it is clear that although there are in principle a sufficient number of TP contacts in the middle regime to determine reasonably accurate structures, it is difficult to find these among the relatively large number of FP contacts. Second, CE-YAPP clearly increases the structural quality also in this regime. Thus, when examining the change in GDT(5) scores (ll*GDT* (5); Fig. 4C) CE-YAPP causes an average increase of 0.12 and 0.09 in the top two ranges of *N*_*Eff*_ */N*_*AA*_. This demonstrates that CE-YAPP is able to improve not only the contact quality but also the structural quality even when there are only an intermediate number of sequences available. Thus, for example, for the 41 proteins in the middle range we find that GDT(5) scores for 32 of the proteins are improved by CE-YAPP.

### Testing other contact prediction methods

The results described above were all obtained using the Gremlin contact predictor [11] to provide the initial set of contacts to CE-YAPP. Contact prediction is, however, a field in rapid development driven both by increases in the number of sequences but also in the ability of improved algorithms [27]. These improvements are having a substantial impact on protein structure prediction, as evident from results from CASP12 [28].

Because CE-YAPP is compatible with any contact predictor we analysed whether the improvements observed are specific to the use of Gremlin, or whether the requirement of structural self-consistency can generically improve a wider range of prediction methods. We thus repeated the contact predictions using four different algorithms, and used these as input to CE-YAPP. Encouragingly we observe a consistent improvement in contact predictions across all methods (Fig. S1).

## Conclusions

We have developed CE-YAPP, a method that automatically chooses a number of (long-range) predicted coevolution contacts as input and increases the precision by removing FP contacts. In its current implementation, CE-YAPP uses secondary structure prediction to define *α*-helical and *β*-stranded segments used to reduce the search space when performing efficient simulated annealing simulations. During the simulations, the weights, *>*_*i*_ (Eq. 3), are sampled to allow systematically-violated restraints to be removed and, thus, identifying them as likely FP contacts. We show, on a selection-bias-free data set consisting of 147 proteins that CE-YAPP increases the precision of PCCs. On average we observe a higher structural quality of the proteins using CE-YAPP contacts.

In the future, we expect the precision of our method should increase synergistically with the development of better contact predictors as well as the addition of system dependent experimental data such as NOEs and/or assigned chemical shift data [22, 24]. We propose CE-YAPP to be used as a fast post-contact-prediction-filter before turning to more advanced structure calculations. Further, it should be possible to combine CE-YAPP with better sampling algorithms and accurate energy functions to obtain improved contact predictions and more accurate three-dimensional structures.

## Methods

### Simulation details

CE-YAPP uses the primary sequence, PCCs implemented as distance restraints, and predicted secondary structure of a target protein as input. Utilizing these sources of information, CE-YAPP performs structure calculations whilst simultaneously identifying and turning off distance restraints that are systematically violated by the 3D geometry. CE-YAPP then performs a final structure calculation with the refined set, keeping the distance restraints fixed. Based on this final structure, the contact list is further refined (Fig. 1). To reduce the noise levels in the refined contact list, the protocol is repeated 64 times and the consensus contacts (*>* 30 % on) are then selected as the final set of contacts.

All simulations were performed using a modified version of the YAPP method [25] implemented in the ALMOST simulation software package [29], and is available at https://sourceforge.net/projects/almost/. Simulated annealing was performed using an implementation of torsion angular dynamics [30, 31] sampling only the dihedrals that are not fixed to ideal secondary structure angles based on a secondary structure prediction. An efficient energy function is used during the simulated annealing that includes a soft-core van der Waals term and a restraints term:

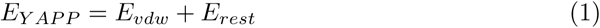

where,

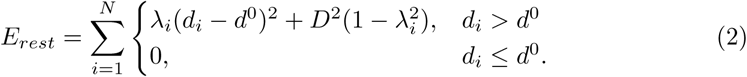

Here, the sum runs over all *N* PCCs, *d*_*i*_ is the C*β*-C*β* (C*α* for GLY) distance in the calculated structure, *d*^0^ = ^7^Å is the distance above which we consider a restraint being violated. *D* is a parameter used to control the acceptable degree of violation, and is decreased during the simulated annealing protocol (see below). The values of *d*^0^, *D* and other key parameters were chosen as described in more detail in the Supporting Information.

During the simulations, the values of *>*_*i*_ (one for each predicted contact) are updated at each time step using a Brownian motion-like equation:

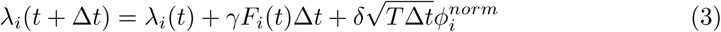

Here *t* is the MD simulation time, ll*t* is the time step, T is the temperature, *F*_*i*_(*t*) is the force exerted by the restraint *i* at a given time *t* and *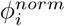* is random noise generated from a standard normal distribution. The parameters *I* and *5* were set to 0.00025 and 0.6666, respectively. All values of *>* were enforced to stay in the range of 0 – 1.

By sampling *>*_*i*_ during the simulations, CE-YAPP can switch specific distance restraints off (*>*_*i*_ = 0) at an energetic cost determined by *D*. During the simulated annealing protocol, *D* is annealed from 150 A^°^ until it reaches 3 Å to steadily remove contacts that are systematically violated. A final structure calculation is performed using the refined list of contacts as fixed restraints (*>*_*i*_ = 1) in a simulated annealing simulation. The restraints that violate the upper limit *d*^0^ by more than *D*, in the final structure, are turned off.

To reduce noise further, we repeat the protocol (Fig. 1) 64 times producing 64 similar contact lists. The repetitions are trivially independent and can, therefore, run simultaneously on a multi-core computer. The contacts that are turned on in more than 30 % of the 64 refined contact lists produced by the 64 repeated protocol runs are then selected for and represent the final contacts produced by CE-YAPP.

In the evaluation of the distance violations of the final set of contacts we also calculated the restraint violation energy:

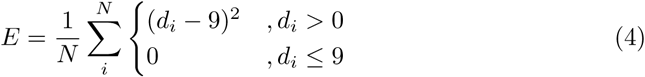

where N is the number of contacts and *d*_*i*_ are the contact distances (CB-CB) observed in the PDB structures (CA for Glycine).

### Fixing the secondary structure

We reduced the conformational space by fixing the secondary structure of simulated proteins to that predicted by PSIPRED [32]. The dihedral angles (*ϕ* and ψ) of the segments predicted to be either *α*-helical or *β*-stranded were fixed to the angles (*ϕ*_*α*_ =*-* 60, *ϕ*_*β*_ = *-*135 and ψ_*α*_ = *-*45, *ψ*_*β*_ = 135) for the respective secondary-structure-type leaving only the remaining regions to change conformation. When using PSIPRED, we refrained from using the default sequence database and used only the MSAs provided with Noumenon to minimize bias that might occur from using the larger default database when predicting secondary structures.

### Effective sequences

The number of effective sequences were calculated by clustering sequences with more than 80 % sequence identity and assigning each sequence within the clusters with a fractional weight of 1*/n*, where *n* is the cluster size. By summing the weights of each sequence one obtains the effective number of sequences which represents the number of diverse sequences in the alignment.

### Contact prediction

We predicted contacts using the stand-alone Gremlin [11] software package using the default settings. Using the predicted secondary structure information, we only select PCCs that do not coincide within a single predicted secondary structure segment, to probe the extraction of long-range contacts. More specifically, we optimised the number of input contacts to be 1.2 times the number of amino acids (*N*_*input*_ = 1.2 *N*_*AA*_). We thus chose this number of contacts among those not found within a fixed secondary structure segment, and used these contacts were then used as distance restraints.

In the analysis of the contacts we define a TP as being a predicted contact with a C*β*-C*β* (C*α* for GLY) distance observed to be at or below 9 Å in the experimental structure.

### Final structure calculations

We performed simulations to determine the structural quality obtained from the different sets of contacts (Fig. 4) using the experimentally-observed dihedral angles extracted from the PDB structures. More specifically, we used STRIDE [33], to determine the secondary structure of the proteins based on the PDB structures, and we extracted the *ϕ* and dihedral angles those residues that were determined (by STRIDE) to be *α*-helical, 3_10_ helical or *β*-stranded. During the simulation, these dihedral angles were kept fixed. In these simulations we also fixed *>*_*i*_ = 1 in Eq. 3 thereby keeping the restraints fixed. We used GDT as a quality measure with a single cut-off of 5 Å. Specifically, we calculated the fraction of C*α* atoms in the structural model that are within 5 Å (GDT(5)) of the corresponding position in the PDB structure. To reduce the noise levels, we take the average GDT(5) of 16 simulations for each set of contacts.

### Computational time

Once the predicted secondary structure (by PSIPRED) and coevolution contacts (from Gremlin) were obtained, the time spent on a single protocol run (Fig. 1) using a single CPU-core (2.3 GHz) takes in the order of 15 CPU-minutes on any of the tested proteins. Thus, with a 64 core machine, the entire protocol can be performed in about 15 mins.

### Protein data

Our benchmark of CE-YAPP was performed using the Noumenon data set [26], which consists of 150 MSAs and their associated protein crystal structures. Three out of the 150 data points were left out of the analysis, simply because their predicted contacts all coincided in unresolved regions of the PDB structures. In particular, when there are only very few effective sequence, Gremlin may score all pairs of columns in the MSA equally, with top ranked contacts then arbitrarily assigned to the N-terminal region. For the three excluded proteins, the N-terminal tails are not resolved in the crystal structures, resulting in data points that we cannot verify against experiments.

## Supporting information

**S1 Fig. Testing on other contact predictors.**

**S2 Fig. Parameter optimization.**

**S3 Fig. Optimising number of input contacts.**

## Acknowledgments

The authors thank Wouter Boomsma for useful discussions and input.

## Other coevolution contact predictors

To ensure that CE-YAPP generalizes across diαerent contact predictors, we applied CE-YAPP on multiple contact predictors. We included plmDCA [1, 2], which, exactly like Gremlin, uses a pseudo-likelihood maximization approach and should be very similar to Gremlin in its predictions. The other methods are, GaussDCA (gDCA [3]), CMAT [4] and PconsC3 [3, 5]. It should be noted, that PconsC3 uses a machine learning approach that combines plmDCA, gDCA and RaptorX [6], for which the latter provides a webserver that we cannot provide the input multiple sequence alignment to. Since, this is a requirement for us, to prevent the selection bias, observed when using too rich multiple sequence alignments, we chose to omit RaptorX and use CMAT as a replacement. We acknowledge that PconsC3 likely underperforms due to this replacement.

We predicted contacts for the Noumenon data set using the above-described methods, and performed contact filtering using CE-YAPP. The precision of the predicted contacts before and after applying CE-YAPP are shown in Fig. S1. Apart from CMAT, the diαerent contact prediction methods perform similarly in terms of precision with CE-YAPP increasing the precision similarly, on the Noumenon data set.

## CE-YAPP parameter selection

Several parameters, in the CE-YAPP method, were either manually tuned or chosen based on previous work [7]. The parameters in question are (I) the number of input contacts, *N*_*input*_, (II) the number of time steps during structure calculations, *N*_*steps*_, (III) the number of repeated simulations, *N*_*repeats*_, and (IV) *D* and *d*^0^ (Eq. 3 in main text).

CE-YAPP was generally robust to changes in the following parameters: *N*_*repeats*_, *N*_*steps*_, *D* and *d*^0^. The values assigned to these parameters were tuned based on the structural accuracy (*Cα*-RMSD) obtained when running

CE-YAPP on three proteins with PDB IDs: 2RQL (95 amino acids), 5P21 (166 amino acids) and 1SVN (269 amino acids). We varied *N*_*repeats*_ and *N*_*steps*_ individually, while *D* and *D*^0^ were varied combinatorially.

Based on the results depicted in Fig. S2, we chose *N*_*steps*_ = 10, 000, *N*_*repeats*_ = 64, *D* = 3 and *d*^0^ = 7. Indeed, for values numbers, the structural accuracy was maximal for the three proteins.

## Number of input contacts

To select the number of input contacts, we varied *X* in

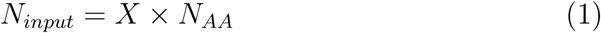

where *N*_*AA*_ is the number of amino acids. We maximized for the total mean structural accuracy across 16 repeated simulations and each of the proteins in the Noumenon data set. In this case, we used the global distance test (GDT) with a single cutoα of 5 Å. In other words, we used a “low resolution” structural accuracy measure that reports on the ratio of *Cα* atoms that are within 5 Å of the experimental PDB structure. As seen in Fig. S3, there is a peak at *X* = 1.2, which we chose as a final parameter. It should be noted that CE-YAPP is fairly robust to changes in *X*, with a mean increase of *∽* 0.04 GDT(5) going from *X* = 0.5 to *X* = 1.2.

---

**Figure S1:**
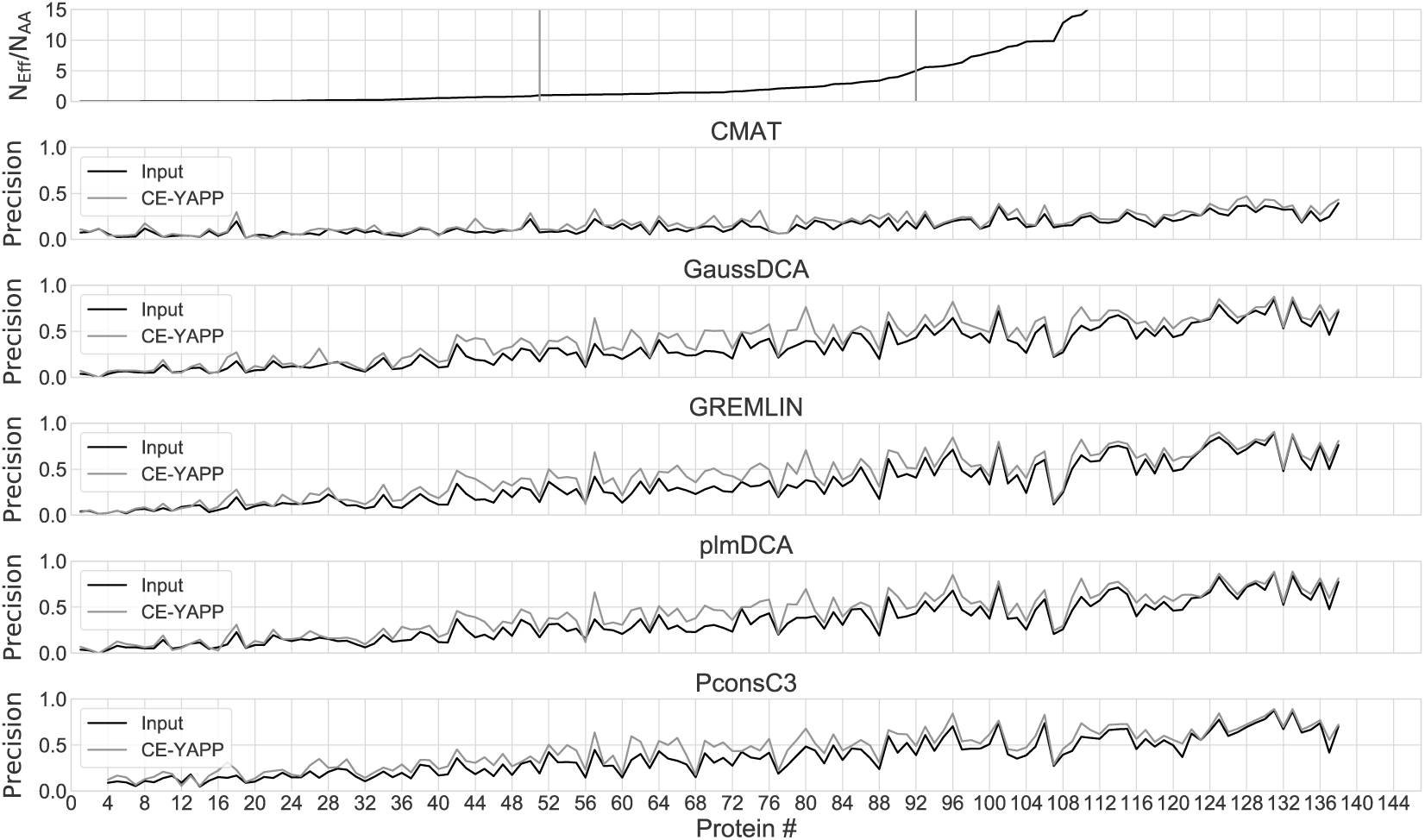
Precision of the Contact Predictors. Panel 1. The number of eαective sequences divided by the number of amino acids, N_Eα_ */*N_AA_, is plotted for each protein, of the Noumenon data set, and sorted from low to high. The data in the remaining panels are sorted accordingly. The grey vertical bars represent the proteins with N_Eα_ */*N_AA_ closest to 1 and 5, respectively. The remaining panels depict the precision (TP/(TP+FP)) of the input contacts and after applying CE-YAPP.

**Figure S2:**
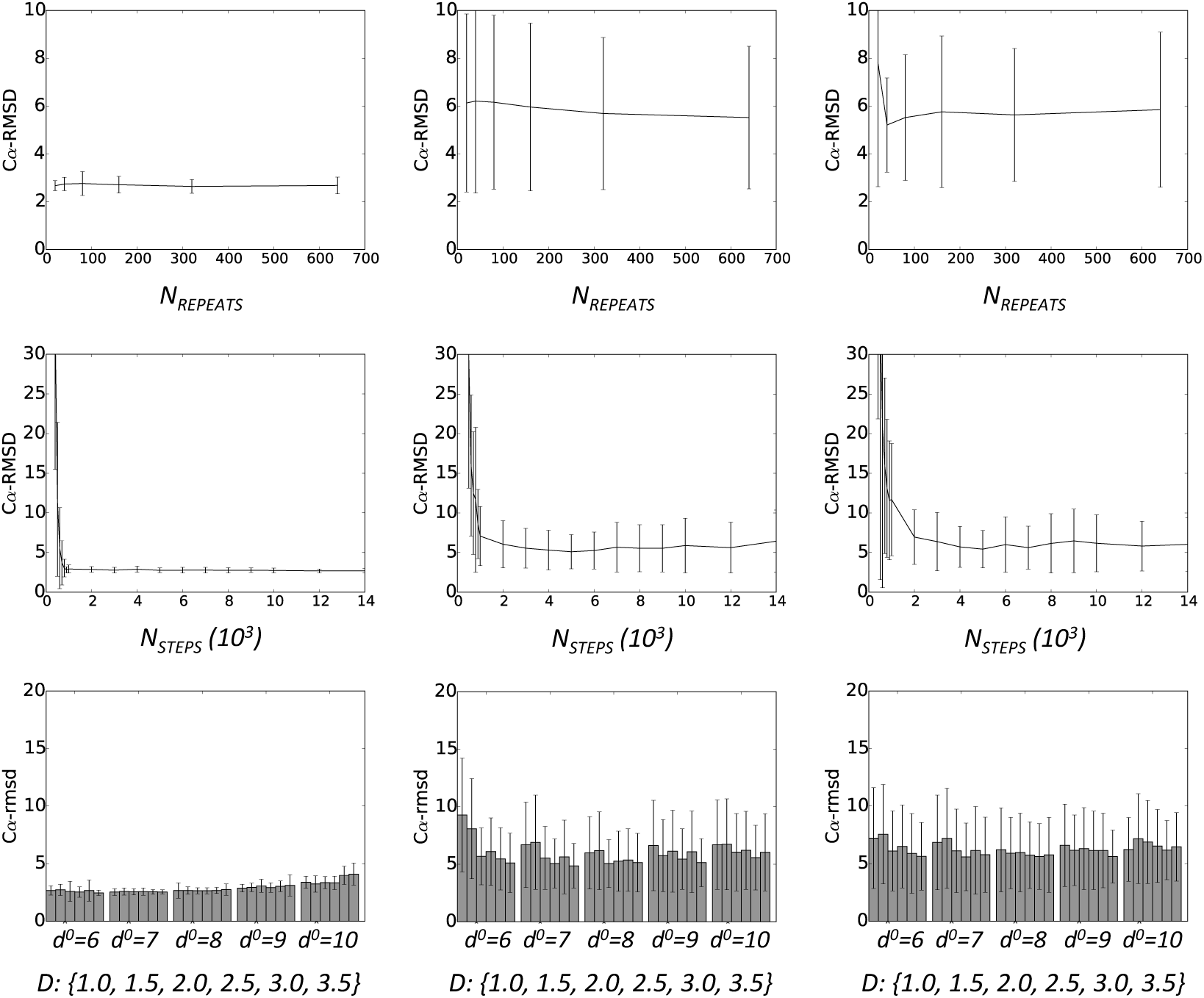
Parameter Sweep. Mean C*α*-rmsd for 50 repeated simulations of three proteins (PDBID: ^2^RQL, ^5^P^21^, ^1^SVN) is plotted with respect to the parameters *Nrepeats* ^(top row), *N*^*steps* ^(middle row), *D*^ and *d*^0^ (bottom row). Each bar plotted in the bottom row represents the mean C*α*-rmsd for 50 repeated simulations with a specific *d*^0^ and *D* (Eq. 3 in main text). AII error bars represent the standard deviation for 50 repeated simulations.

**Figure S3:**
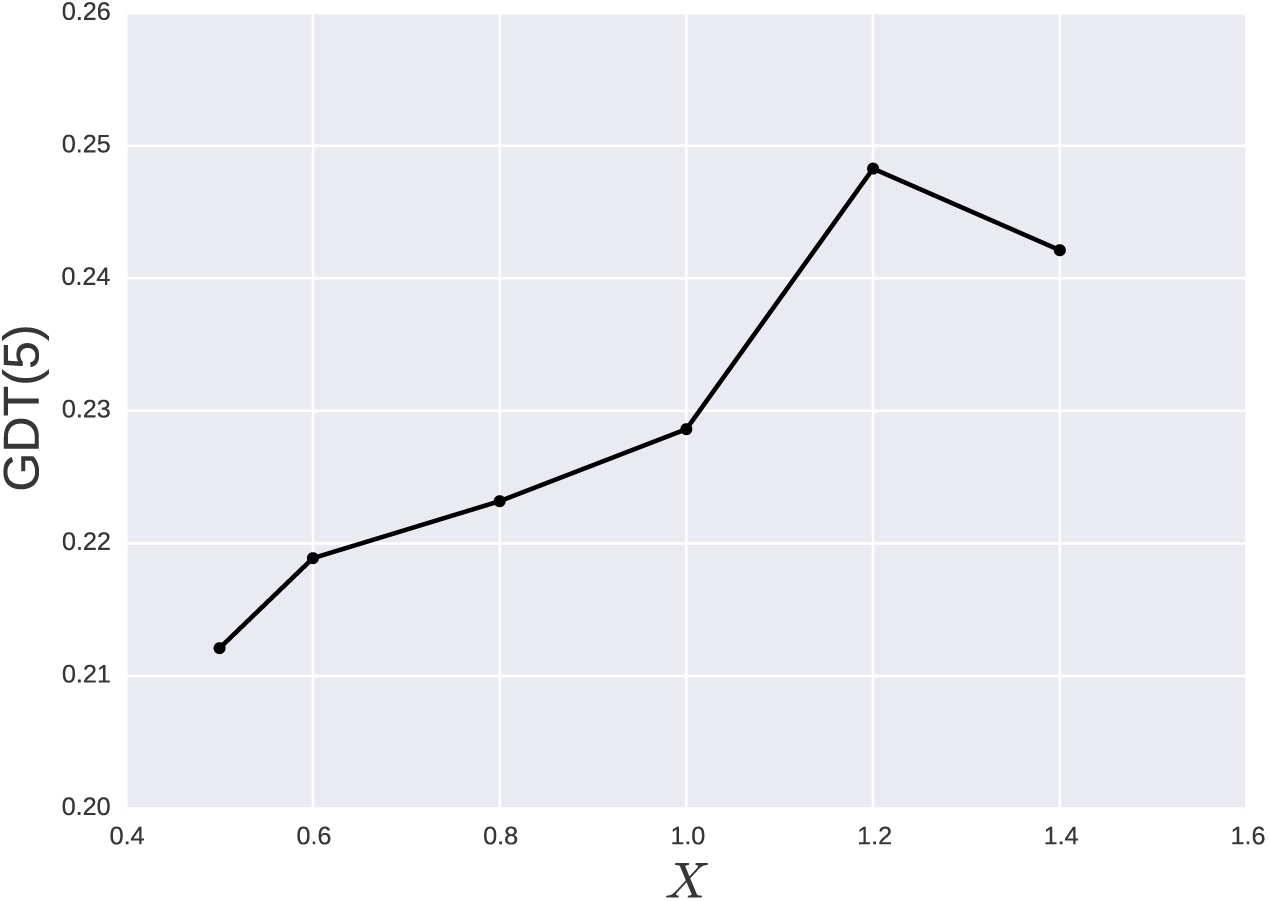
Number of Input Contacts. Total mean GDT(5) across 16 repeated simulations and each of the proteins in the Noumenon data set is plotted as a function of *X*, described in Eq. 1 in the Supporting Information.

